# Molecular signatures of cognition and affect

**DOI:** 10.1101/2020.07.16.203026

**Authors:** Justine Y. Hansen, Ross D. Markello, Jacob W. Vogel, Jakob Seidlitz, Danilo Bzdok, Bratislav Misic

## Abstract

Regulation of gene expression drives protein interactions that govern synaptic wiring and neuronal activity. The resulting coordinated activity among neuronal populations supports complex psychological processes, yet how gene expression shapes cognition and emotion remains unknown. Here we directly bridge the microscale and macroscale by mapping gene expression patterns to functional activation patterns across the cortical sheet. Applying unsupervised learning to the Allen Human Brain Atlas and Neurosynth databases, we identify a ventromedial-dorsolateral gradient of gene assemblies that separate affective and cognitive domains. This topographic molecular-psychological signature reflects the hierarchical organization of the neocortex, including systematic variations in cell type, myeloarchitecture, laminar differentiation, and intrinsic network affiliation. In addition, this molecular-psychological signature is related to individual differences in cognitive performance, strengthens over neurodevelopment, and can be replicated in two independent repositories. Collectively, our results reveal spatially covarying transcriptomic and cognitive architectures, highlighting the influence that molecular mechanisms exert on psychological processes.

## INTRODUCTION

The human brain is an integrated system, involving interactions across multiple scales [9, 50]. At the molecular level, fluctuations in gene expression and protein synthesis in neurons drive single-cell activity [16, 66, 70]. The waxing and waning of cellular activity promotes synaptic remodeling [30, 48, 84], shaping the wiring of nested and increasingly polyfunctional neural circuits [11, 17, 62]. Anatomical connections among mesoscopic neuronal populations promote functional interactions [69], manifesting as patterned neural activity that drives psychological processes [6, 21, 68, 87]. The regulation of gene expression is therefore naturally intertwined with the brain’s structure and function [10, 32, 33, 64, 76]. How molecular dynamics map onto mental states remains a key question in neuroscience [8].

Modern technological and analytic advances, in concert with global data-sharing initiatives, have created fundamentally new opportunities to link molecular dynamics and psychological processes. High-resolution functional neuroimaging has informed comprehensive meta-analytic atlases of how brain areas selectively respond over a spectrum of perceptual, cognitive and affective experimental manipulations [12, 24, 29, 34, 37, 51, 86]. At the same time, high-throughput microarray profiling has yielded precise genome-wide maps of transcript distributions over the brain [39, 40, 58], allowing inferences about the spatial distribution of cellular processes and types [3, 4, 14, 32, 33, 64, 65, 67, 74, 83]. Altogether, the concurrent emergence of global functional genomic and brain mapping initiatives offers an unprecedented chance to identify spatial correspondences be-tween the brain’s genetic and cognitive architectures.

Here we directly relate microscale molecular processes to the macroscale functional architecture of the human brain. We apply partial least squares analysis to gene expression maps (Allen Human Brain Atlas; [40]) and probabilistic functional activation maps (Neurosynth; [86]) to identify molecular signatures related to psychological processes (for a conceptually similar approach, see [27]). We reveal distinct sets of functionally inter-related genes that underlie a cognitive-affect gradient of functional processes, and show that this molecular signature corresponds to systematic variation in cell type compositions, microstructure, and large scale functional system affiliation. Using data from the Human Connectome Project (HCP; [72]) we also show that this gene expression signature mediates individual differences in behaviour via microstructure. Finally, we perform extensive cross-validation, sensitivity testing and replication using two independent datasets (BrainMap; [29], and BrainSpan; [58]).

## RESULTS

To establish a relationship between gene expression and functional activity, we used the Allen Human Brain Atlas for estimates of gene expression in the brain [40], and Neurosynth for probabilistic measures that specific terms (such as “attention”, “emotion”, and “sleep”) are functionally related to specific brain regions [86]. This probability describes how often specific terms and voxel coordinates are published in conjunction with one another. To facilitate comparison with other reports, only genes with a differential stability greater than 0.1 were retained for analysis (see *Methods*; [14, 39]), and the term set was restricted to those in the intersection of terms reported in Neurosynth and in the Cognitive Atlas [63]. Gene expression data and probabilistic measures were parcellated into 111 left hemisphere cortical regions of interest [18, 23]. The resulting gene expression matrix was composed of normalized expression levels of 8825 stable genes across 111 target brain regions [18], and the functional activation matrix represented the functional relatedness of 123 terms to the same 111 brain regions.

### Molecular signatures of psychological processes

We related gene expression to functional activation using partial least squares analysis (PLS), a multivariate statistical technique that extracts optimally covarying patterns from two data domains [45, 53, 54, 85] (Fig. 1a). PLS analysis revealed a single statistically significant latent variable relating gene expression to corresponding functional activation across the brain (*p*_spin_ = 0.0228), where significance was assessed using a permutation test that preserves spatial autocorrelation (“spin test”) [2, 75]. This latent variable represents a pattern of gene expression (gene weights) and a pattern of functional activation (term weights), that together capture 65% of the covariance between gene expression and functional activation (Fig. 1b). Projecting the gene expression and functional activation matrices back onto the gene weights and term weights, respectively, reflects how well a brain area exhibits the gene and term pattern, which we refer to as “gene scores” and “term scores” (Fig. 1c). The pattern of gene and term scores across the brain revealed a dorsolateral to ventromedial gradient, in which dorsolateral regions were scored more negatively and ventromedial regions more positively.

**Figure 1.**
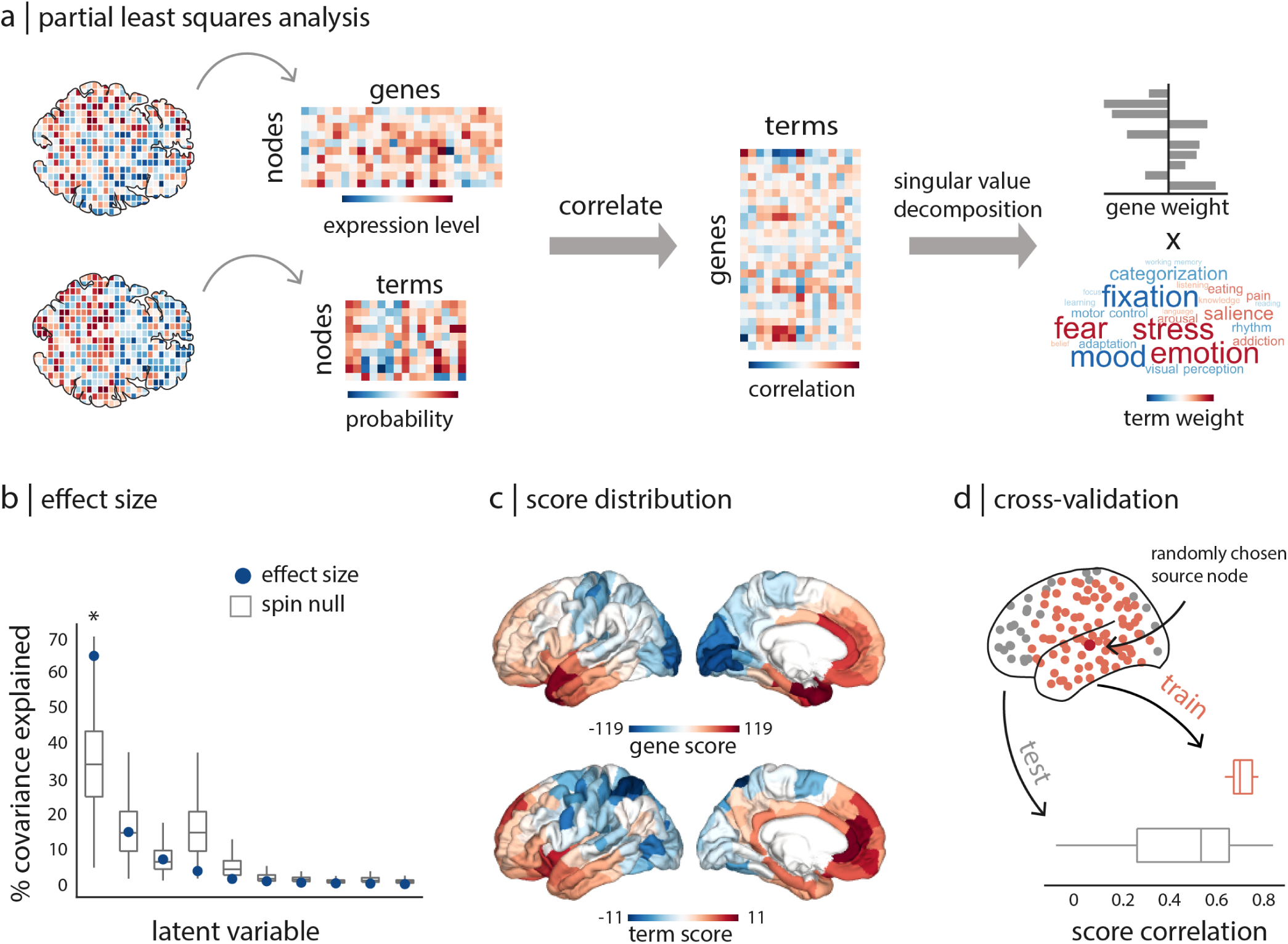
Relating gene expression to functional activation. Partial least squares analysis (PLS) was used to identify spatially covarying patterns of gene expression (Allen Human Brain Atlas) and functional activation (Neurosynth). (a) PLS relates two data domains by correlating the variables across brain regions and subjecting this to singular value decomposition. This results in multiple latent variables: linear weighted combinations of the original variables (gene weights and term weights) that maximally covary with each other. (b) Latent variables are ordered according to effect size (the proportion of covariance explained between gene expression and functional activation they account for) and shown in blue dots. Statistical significance is assessed with respect to spatial autocorrelation-preserving null model [2], shown in grey. Only the first latent variable was statistically significant (*p*_spin_ = 0.0228), accounting for 65% of the covariance between gene expression and functional activation. (c) Projecting the original data back onto the PLS-defined gene/term weights results in gene/term scores for each brain region, indexing the extent to which a brain region expresses covarying gene/term patterns. (d) The correlation between gene scores and term scores was cross-validated by constructing the training set with 75% of brain regions closest in Euclidean distance to a randomly chosen source node (red), and the testing set as the remaining 25% of brain regions (grey). See Fig. S1 for a comparison with completely random splits.

We next cross-validated the correlation between gene and term scores. Due to inherent spatial autocorrelation, proximal regions exhibit similar gene expression profiles and functional activity [2, 15]. Thus, randomly dividing brain regions into training and testing sets may result in interdependencies between the two sets (Fig. S1). To ensure that the correlation between gene and term scores is not inflated due to spatial autocorrelation, we selected the 75% of brain regions closest in Euclidean distance to a randomly chosen source node as the training set, and the remaining 25% of brain regions as the testing set. This procedure was repeated 100 times and a distribution of correlations for the training and testing set is shown in Fig. 1d. The mean out-of-sample correlation between gene and term scores was 0.4770.

### Distinct gene assemblies underlie cognition and affect

The significance of the first latent variable and the cross-validation of score correlations demonstrates there is a robust relationship between gene expression and functional activation. The relationship itself is determined by the terms and genes that contribute most to the latent variable. The loading of each term was computed as the correlation between the term’s functional activation across brain regions with the PLS-estimated scores. The 25% most positively and negatively correlated terms were retained as terms that most contribute to the latent variable (Fig. 2a; for the loadings of all reliable terms, see Fig. S2). Terms with large positive loadings were related to affective processes, including emotion, stress, fear, anxiety, and mood. Terms with large negative loading were identified as terms related to higher-order cognitive processes. Examples include attention (of which “visual attention”, “spatial attention”, and simply “attention” were all weighed very highly), visual perception, and imagery. This latent variable thus represents a putative cognitive-affective gradient of functional activity.

**Figure 2.**
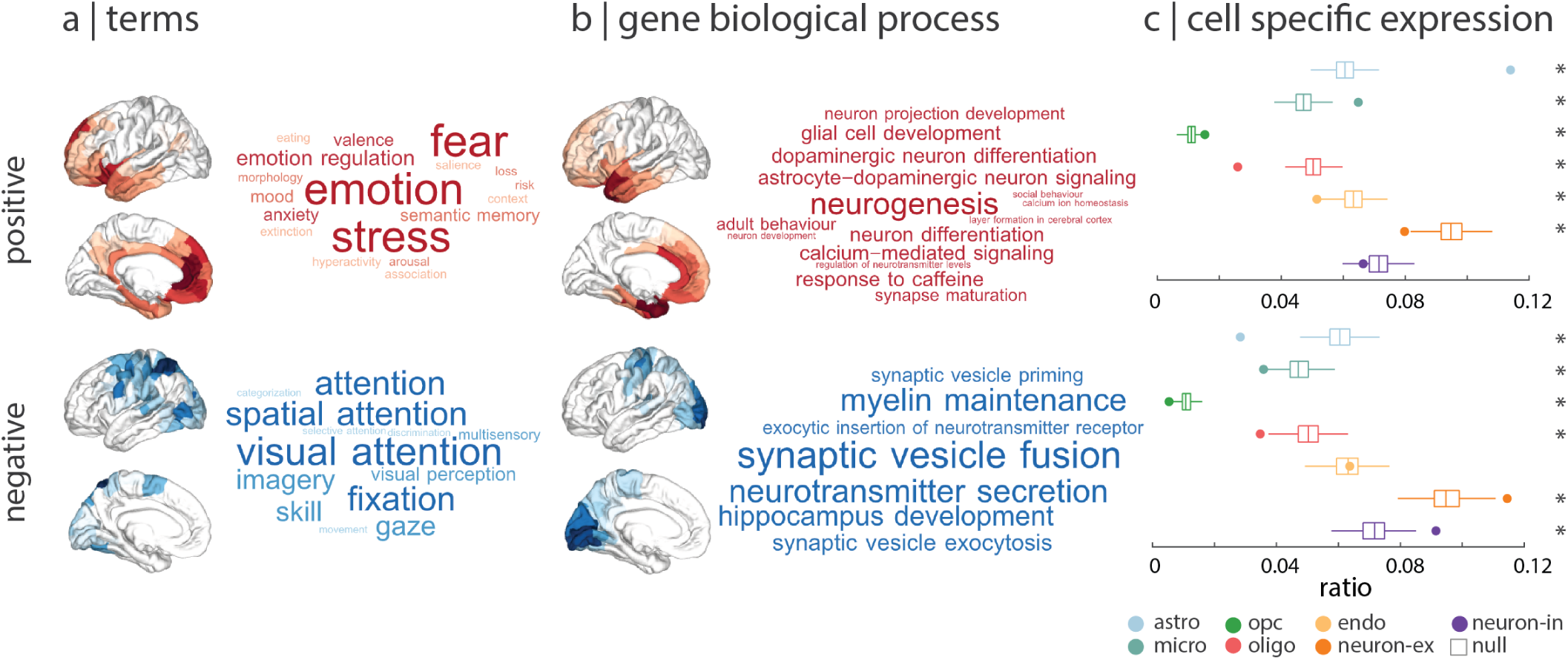
Gene sets underlying cognition and affect. Genes and terms that contribute most to the latent variable were analyzed further. (a) The contribution of positively- and negatively-weighed ontological terms to the latent variable were estimated with term loadings: correlations between a term’s functional activation across brain regions and the PLS-derived score pattern. The 25% most positively (red) and negatively (blue) correlated terms demonstrate a cognitive-affective gradient. Word size represents the relative size of the term loading. (b) Gene contribution was estimated with gene loadings: correlations between a gene’s expression across brain regions and the PLS-derived score pattern. Biological processes in which the top 50% of genes with positive and negative loading are most involved were identified using Gene Set Enrichment Analysis (see *Methods*) and tested against a spatial autocorrelation-preserving null model [31]. (c) Cell-type deconvolution was used to identify cell type enrichment in the gene sets identified by PLS [67]. The ratio of genes in each gene set preferentially expressed in seven distinct cell types is shown against a null model of a random selection of all genes (boxplots). Cell types: ASTRO = astrocyte, MICRO = microglia, OPC = oligodendrocyte precursor, OLIGO = oligodendrocyte, ENDO = endothelial, NEURO-EX = excitatory neurons, NEURO-IN = inhibitory neurons, NULL = empirically derived null distribution.

Gene contribution was analogously assessed by computing spatial correlations (loadings) between an individual gene’s expression pattern and the PLS-derived scores. Genes with the 50% most positive loadings (which we subsequently refer to as “positive genes”) covary with the functional activity of terms related to affective processes, and genes with the 50% most negative loadings (“negative genes”) covary with the functional activity of terms related to higher-order cognitive processes. In other words, the pattern of positive genes covarying with affective terms is strongest in positively scored brain regions (Fig. 2a), and the pattern of negative genes covarying with cognitive terms is strongest in negatively scored brain regions (Fig. 2b).

To better understand the biological significance of the positive and negative gene sets, we adapted analyses from the Gene Set Enrichment Analysis toolbox (https://github.com/benfulcher/GeneSetEnrichmentAnalysis [31]). We explored the biological processes with which the reliable positive and negative genes are significantly involved (see *Methods* for details and Table S4 and S5 for a full machine-readable list of biological processes and respective *p*-values). A selection of the significant categories most related to brain structure and function are visualized as word clouds in Fig. 2b. In general, affect-related gene sets show enrichment for processes related to neurogenesis and differentiation, while cognition-related gene sets are enriched for processes related to synaptic signaling.

Alongside biological process, we asked whether psychologically-relevant genes are preferentially expressed in specific cell types (Fig. 2c). Cell-type deconvolution was performed using cell-specific aggregate gene sets across five human adult postmortem single-cell and single-nucleus RNA sequencing studies ([22, 38, 47, 49, 55, 88]), as presented previously (Supplementary Table 5 from [67]). Specifically, we calculated the ratio of genes in each gene set preferentially expressed in one of seven cell types: astrocytes, microglia, oligodendrocyte precursors, oligodendrocytes, endothelial cells, excitatory neurons, and inhibitory neurons (Fig. 2c). Gene sets were thresholded to include the top 50% of genes with greatest loadings (note that although the threshold is arbitrary, the results are highly consistent across a range of thresholds, from 2.5% to no threshold; Fig. S3). Statistical significance was assessed against a null distribution of ratios constructed by repeating the process 10 000 times on a set of random genes (two-tailed, FDR-corrected). Dominant positive genes (related to affect) are significantly more expressed in astrocytes (*p* = 2.3 *×* 10^−4^), microglia (*p* = 2.3 *×* 10^−4^), and oligodendrocyte precursors (*p* = 0.0160), and significantly less expressed in excitatory neurons (*p* = 0.0052), oligodendrocytes (*p* = 2.3 *×* 10^−4^), and endothelial cells (*p* = 0.0052). Dominant negative genes (related to cognition) are significantly more expressed in excitatory neurons (*p* = 0.0023) and inhibitory neurons (*p* = 0.0017), and significantly less expressed in astrocytes (*p* = 0.0007), microglia (*p* = 0.0114), oligodendrocytes (*p* = 0.0019) and oligodendrocyte precursors (*p* = 0.0114). Broadly, we find evidence that areas associated with affect are enriched for genetic signal of cells involved in neuron support (astrocytes, microglia); areas associated with cognition are enriched for genetic signal of neurons themselves (inhibitory and excitatory). This dichotomy also matches the intuition derived from biological process enrichment analysis (Fig. 2b).

### The gene-activation gradient is organized around microscale and macroscale hierarchies

Having identified a gradient of covarying gene expression and functional activation, we next investigated whether these topographic patterns reflect variation in other microstructural and functional attributes [42, 81]. To address this question, we averaged measures of cortical thickness and T1w/T2w ratios (a widely used proxy for intracortical myelin; [36]) from the left hemisphere cortex across 417 unrelated subjects from the Human Connectome Project (see *Methods*). We then computed Pearson’s correlations of mean cortical thickness and T1w/T2w maps with gene score and term score maps (Fig. 3a). We find a strong positive correlation between cortical thickness and PLS scores (*r* = 0.8216, *p*_spin_ = 0.0002 for gene scores, *r* = − 0.5201, *p*_spin_ = 0.0426 for term scores), and a strong negative correlation between T1w/T2w ratio and PLS scores (*r* = − 0.8586, *p*_spin_ = 0.0003 for gene scores, *r* = − 0.5822, *p*_spin_ = 0.0039 for term scores). Altogether, the gene-activation gradient mirrors microstructural attributes [14, 33, 41, 80].

**Figure 3.**
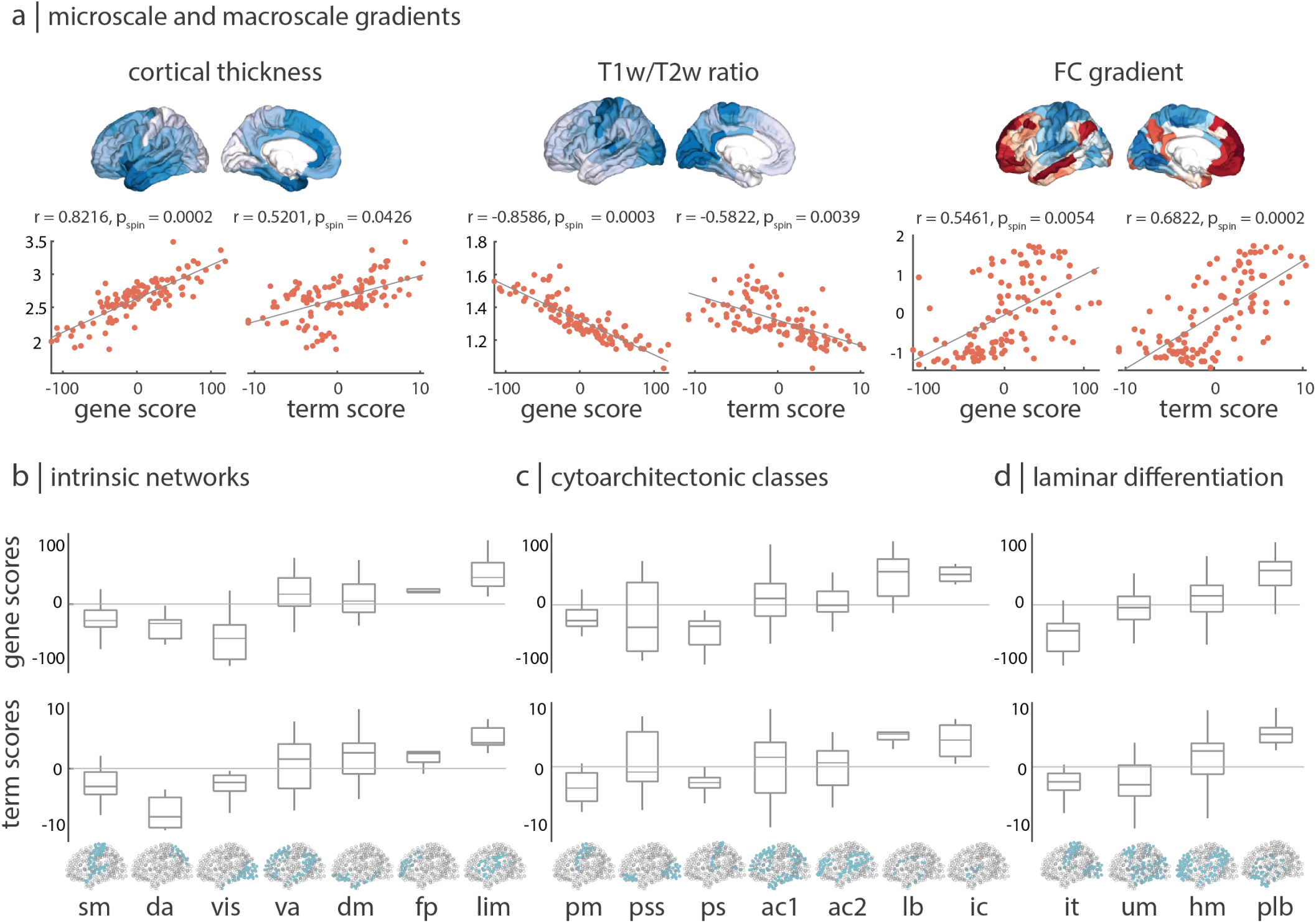
The gene expression-functional activation gradient is organized around microscale and macroscale hierarchies. (a) PLS-derived score patterns are positively correlated with cortical thickness, negatively correlated with intracortical myelin (measured by T1w/T2w ratio), and positively correlated with the principal gradient of functional connectivity. (b) Distribution of scores across seven intrinsic resting-state functional networks defined by Yeo and colleagues [87]. (c) Distribution of scores across the seven Von Economo cytoarchitectonic classes [76, 78, 79]. (d) Distribution of PLS-derived gene and term scores across the four Mesulam levels of laminar differentiation [57, 60]. Network assignments: SM = somatomotor, DA = dorsal attention, VIS = visual, VA = ventral attention, DM = default mode, FP = fronto-parietal, LIM = limbic, PM = primary motor cortex, PSS = primary/secondary sensory cortex, PS = primary sensory cortex, AC1, AC2 = association cortex, LB = limbic regions, IC = insular cortex, IT = idiotypic, UM = unimodal, HM = heteromodal, PLB = paralimbic.

Given that the score pattern resembles the differentiation between unimodal and transmodal cortex [43, 56], we sought to relate the score pattern to the principal functional gradient reported by Margulies and colleagues [52]. For this purpose, we applied diffusion map embedding on a group-averaged functional connectivity matrix computed from the 1 003 HCP subjects with complete resting-state fMRI data, and extracted the first principal gradient [19, 71]. This gradient situates brain regions on a continuous axis from unimodal primary sensory and motor cortex to transmodal higher association cortex (Fig. 3a). We find that the gene score and term score patterns significantly correlate with this gradient (*r* = 0.5461, *p*_spin_ = 0.0054 for gene scores, *r* = 0.6822, *p*_spin_ = 0.0002 for term score; Fig. 3a). This implies that negatively scored regions tend to be more closely aligned with unimodal cortex and positively scored regions tend to be predominantly aligned transmodal cortex.

As a final step, we sought to understand how well the gene and term score maps conform to other major structural and functional partitions of the human cerebral cortex. We stratified gene and term scores in several complementary ways: (1) within seven intrinsic functional brain networks as defined by Yeo and colleagues [87], (2) within seven Von Economo classes of cortical cytoarchitecture [76, 78, 79], and (3) within four Mesulam levels of laminar differentiation across the cortex [57, 60] (Fig. 3b–d). Consistent with the notion that the gene expression-functional activation gradient reflects a differentiation between cognitive and affective psychological domains, we observe a separation between limbic/paralimbic and somato-motor/idiotypic networks across all three partitions.

### Relating psychologically-informed patterns of gene expression to individual differences in behaviour

Finally, we asked whether a physical manifestation of topographic PLS score maps is related to interindividual performance differences on established psychometric assessments of cognition and affect. Since PLS-derived gene scores are most highly correlated with T1w/T2w ratios (Fig. 3a), we used T1w/T2w ratios from *n* = 417 unrelated HCP subjects as a proxy for the degree to which the gene expression-functional activation gradient is observed in a given individual. Specifically, we correlated individual subjects’ T1w/T2w ratios with the PLS gene scores (Fig. 4a). This approach results in a distribution of subject correlations that describe how well an individual manifests the gene score pattern by means of their T1w/T2w ratio. This vector of correlations per subject was then independently correlated to the performance on 58 behavioural measures (see *Methods* for details of variable selection and Table S3 for a complete list of behavioural measures included). These measures include specific psychological tests (i.e. the Pennsylvania Matrix Test for fluid intelligence), composite scores (i.e. a composite score of executive cognitive function), and individual subjective rankings of emotional states and social traits (i.e. rating how stressed one feels or how many close friends they have). The procedure yielded a correlation for each behavioural measure, indicating whether there is a relationship between how well an individual physically manifests the PLS gene score pattern and their score on the behavioural assessment (Fig. 4b–c). Significance was determined using an FDR corrected two-tailed permutation test in which the null distribution was constructed by correlating the original behavioural performances to a permuted vector of the correlations between gene scores and T1w/T2w ratios [7].

**Figure 4.**
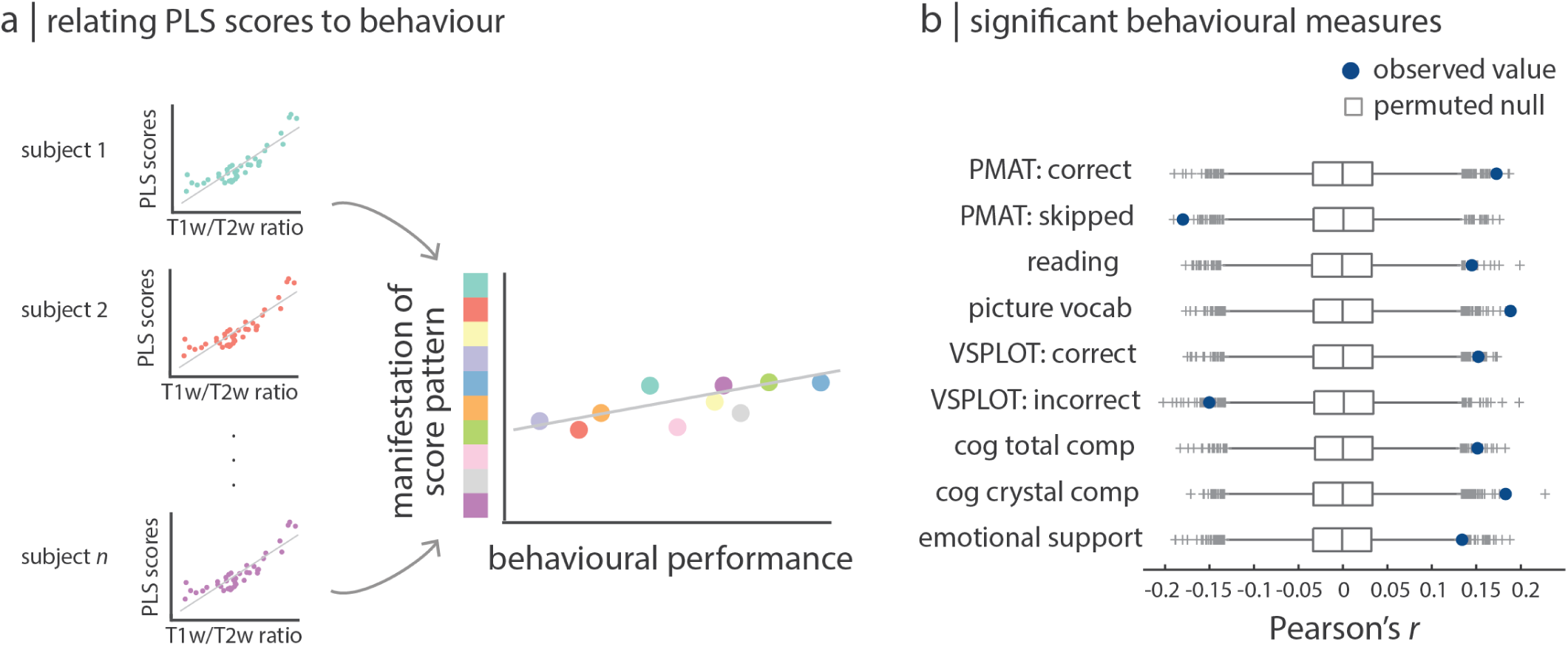
Molecular signatures of cognition influence individual differences in behaviour. We asked whether functionally-relevant patterns of gene expression mediate individual differences in behaviour via brain structure, using T1w/T2w maps and behavioural assessments from *n* = 417 unrelated HCP subjects. (a) For each subject, T1w/T2w maps were independently correlated to the PLS-derived gene score pattern. This correlation represents the degree to which a subject exhibits the gene score pattern (Fig. 1c). These individual-specific correlation coefficients were then correlated with individual performance on a range of 58 behavioural assessments. (b) Subjects who best manifest the gene score pattern via their T1w/T2w ratio performed significantly better on eight cognitive tasks and one subjectively scored measure. Significance was assessed using an FDR corrected two-tailed permuted test, where null models are shown as grey boxplots. For visualization, gene scores were multiplied by − 1 to give positive correlations with better performance. Boxplots represent the 1st, 2nd (median) and 3rd quartiles, whiskers represent the non-outlier end-points of the distribution, and crosses represent outliers.

We find that individuals whose T1w/T2w map is more similar to the negative gene score pattern perform better on behavioural tests of cognitive function. Specifically, the significantly correlated cognitive tests included: the number of correct responses on the Pennsylvania Matrix Test (a measure of fluid intelligence, *r* = 0.1727, *p* = 0.0130), the Reading Test (a measure of reading decoding skills and crystallized cognitive abilities, *r* = 0.1450, *p* = 0.0232), the Picture Vocabulary Test (a measure of general vocabulary knowledge and crystallized cognitive abilities, *r* = 0.1885, *p* = 0.0039), the number of correct responses on the Short Pennsylvania Line Orientation Test (a measure of spatial orientation, *r* = 0.1523, *p* = 0.0232), the Cognitive Function composite score (a composite score of fluid and crystallized cognitive ability, *r* = 0.1513, *p* = 0.0232), and the Crystallized Cognitive composite score (a composite score of of verbal reasoning, *r* = 0.1829, *p* = 0.0039). In addition to these cognitive tasks, a significant correlation was also observed for subjectively scored emotional support (*r* = 0.1340, *p* = 0.0432). Altogether, we find that functionally-relevant patterns of gene expression may mediate individual differences in behaviour via brain structure.

### Molecular signature of psychological function strengthens with development

Given the continuous development of cognitive processes over the lifespan, we sought to track the gene expression-functional activation signature through human development. We used BrainSpan, a dataset that provides gene expression estimates from brain tissue samples aged eight post-conception weeks (pcw) to forty years, across sixteen unique cortical regions. A gene expression matrix was constructed for five different life stages: fetal, infant, child, adolescent, and adult (see *Methods* for details). Gene scores for the twelve regions with available gene expression estimates across all five life stages were estimated by projecting each gene expression matrix onto the PLS-derived gene weights. This results in a set of scores for twelve regions across brain development (Fig. 5a). The molecular signature, represented by gene scores, increases with development, suggesting the gene expression-functional activation gradient becomes more pronounced with maturation. In other words, the genetic signal captured by the original PLS analysis is specific to adult-derived cells.

**Figure 5.**
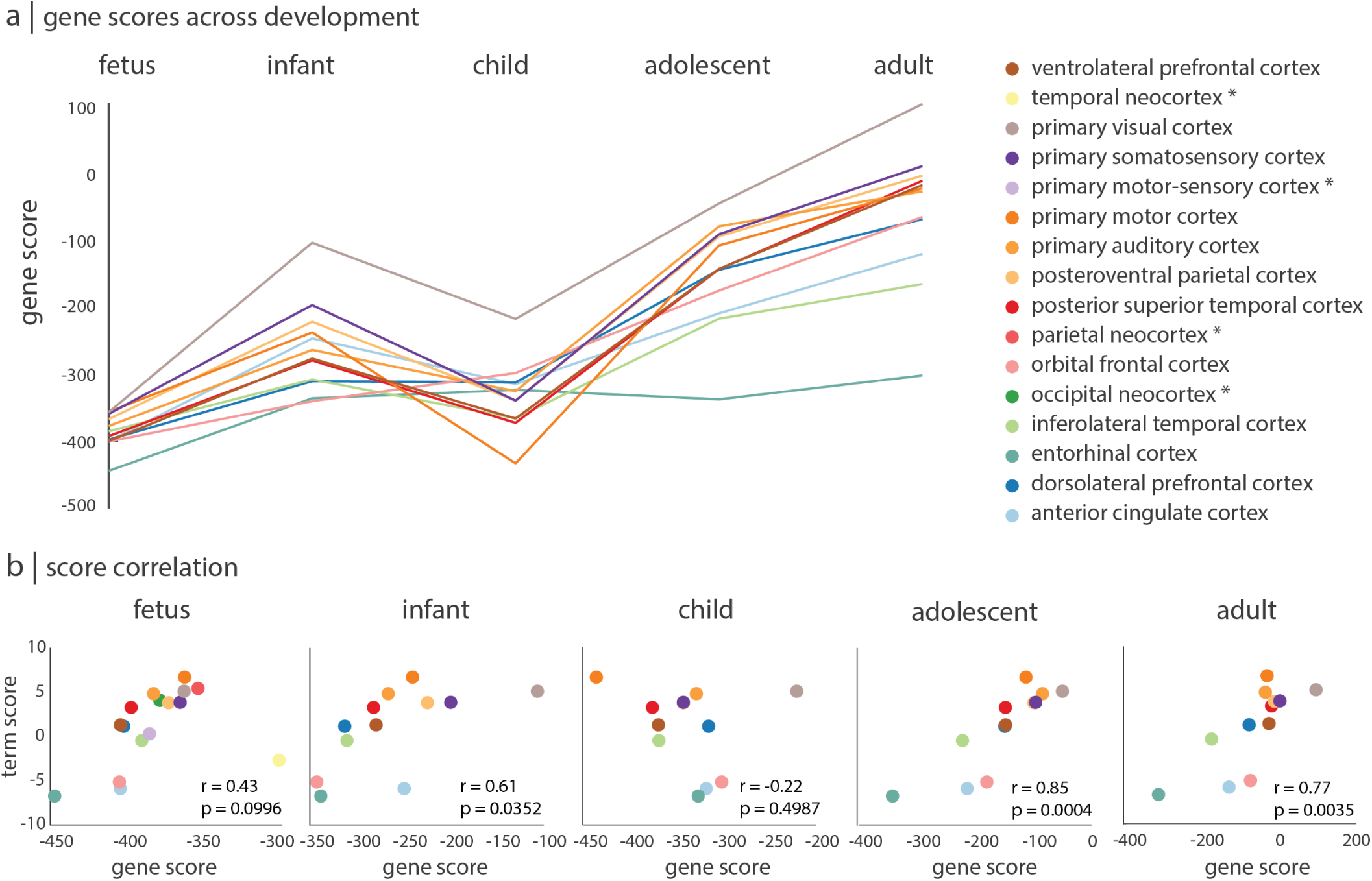
Molecular signature of psychological function strengthens with development. The BrainSpan database was used to replicate the results and also to compare how the isolated genetic signature develops over the lifespan [58]. (a) Gene scores for twelve unique brain regions gradually increase with development and peak in adulthood. Brain regions with gene expression levels only available in the fetal stage are indicated with an asterisk, and no corresponding curve is shown. (b) The correlation between estimated gene scores and PLS-derived term scores is strongest in adolescence and adulthood.

We also used the BrainSpan dataset to externally validate the original PLS model. Specifically, we mapped the 16 unique cortical regions to the 34-node parcellation and averaged PLS-derived term scores from the 34-node parcellation across sibling nodes relating to a parent node. This allows us to correlate estimated gene scores with term scores at each stage of development (Fig. 5b). Estimated gene scores and term scores correlated significantly in the infant (*r* = 0.61, *p* = 0.0352), adolescent (*r* = 0.85, *p* = 0.0004), and adult (*r* = 0.77, *p* = 0.0035) life stages.

### Sensitivity and validation analysis

All analyses presented thus far were conducted on a particular parcellation of brain regions and a predefined set of genes. To ensure the observed results are not dependent on these methodological choices, we compared results when analyses were repeated across different node resolutions and gene sets. Furthermore, we replicated the results using a second dataset for the construction of the functional activation matrix.

To ascertain our findings against different choices of parcellation resolution, gene expression and functional activation matrices were parcellated into three resolutions: a 34-node parcellation, a 57-node parcellation, and a 111-node parcellation [18]. Importantly, the 57-node and 111-node parcellations are derived by dividing the 34-node parcellation into smaller parcels, such that each node in the 57- and 111-node parcellations is a child of a node in the 34-node parcellation. Furthermore, since the original analyses were conducted on differentially stable genes (see *Methods*), and the calculation of differential stability depends on parcellation resolution, the number of stable genes retained across parcellations varied (between 8 825 and 11 560 genes retained). The gene expression and functional activation matrices at these three resolutions were subjected to PLS and the gene scores and term scores were computed. In the two finer resolutions, the mean score of sibling nodes was computed such that one score for each of the 34 parent nodes was available for all three resolutions. Correlating gene scores and term scores across resolutions revealed an almost one-to-one relationship, indicating node resolution has little impact on gene scores and term scores (Fig S4).

Likewise, we asked whether the gene sets contributing most to the latent variable would be altered based on which genes were included in the gene expression matrix. Since the two gene sets underlying cognition and affect are defined based on gene loadings, we compared loadings of genes across six variations of the gene expression matrix. For each of the three resolutions introduced above, one matrix includes all 20 323 genes and another includes only differentially stable genes, as defined by the specific parcellation. Each of the six gene expression matrices, alongside their corresponding functional activity matrix, were subjected to PLS analysis and loadings were computed for each gene, where genes with the top 50% of positive and negative loadings are considered reliable. When compared across the six different gene expression matrices, we find that reliable gene sets are highly consistent (Fig. S5).

We next replicated original results using a different data source for the construction of the functional activation matrix. BrainMap is a manually curated database of published voxel coordinates from neuroimaging studies that are significantly activated or deactivated during tasks [28, 29, 46, 73]. Using the analytic pipeline we previously applied to Neurosynth, we converted Brain-Map data into a functional activation matrix of probabilities, which included 66 terms (see *Methods* for details, and Table S2 for a full list of terms).

PLS analysis on the original gene expression matrix with this BrainMap-derived functional activation matrix again revealed a single statistically significance latent variable that captured 51% of the covariance between gene expression and functional activation (*p*_spin_ = 0.0034). The gene and term score distributions again follow a ventromedial-dorsolateral gradient (Fig. S6a), and gene weights were highly correlated with the original Neurosynth-derived gene weights (Fig. S6b). Term loadings were computed and the reliable positively and negatively correlated terms are shown in Figure S6c. Unlike the terms used in the Neurosynth-derived functional activation matrix, some terms in BrainMap were pharmacological in nature. Interestingly, positive pharmacology terms are primarily depressants (like alcohol and marijuana), and negative pharmacology terms are primarily stimulants (like caffeine).

## DISCUSSION

In the present report, we identify spatially covarying gradients of gene expression and functional activation across the neocortex. Collectively, these patterns delineate a ventromedial-dorsolateral axis, separating gene sets related to cognitive versus affective function. The spatial patterning of gene and term scores follows a hierarchical organization, is closely related to multiple structural and functional attributes, and to individual differences in behaviour. We externally validate our results in two distinct datasets and show that the gene-activation signature strengthens with human development. Our results directly bridge microscale gene expression to macroscale functional processes and highlight the influence that molecular mechanisms have on cognition and behaviour.

The present findings build on previous reports that link gene expression to the structural and functional architecture of the brain. Gene expression profiles have been linked to cortical folding [1], cortical shrinkage during adolescence [83], subcortical connectivity [26], and patterns of long-distance and short-distance neural communication [32, 44, 76]. In particular, intracortical myelin distribution, as measured by T1w/T2w ratio, is correlated with regional transcription levels, potentially reflecting a hierarchical axis of cytological properties, including cytoarchitecture and cell density [14, 33]. Our results expand on this literature, demonstrating that gradients of gene expression distinguish affective processes from cognitive processes. In other words, by shaping micro- and macro-scale brain structure, gene expression is naturally related to neurocognitive organization and, ultimately, to psychological function [27].

What do the present findings show us about the regional specialization for psychological functions? Although we have summarized the gene-activation gradient as one primarily differentiating cognitive and affective processes, greater nuance is warranted. In particular, the posterior/dorsal system is more specifically related to perception, orienting and attention, whereas the anterior/ventral system is more specifically related to emotion and evaluation. Thus, the axis differentiates attentional and evaluation functions, and may be more aptly termed an “affective-attentive” or “evaluation-perception” axis. Interestingly, many of the intermediate terms that do not load highly on either end of the axis are integrative in nature (e.g. “consciousness’, “integration”, “episodic memory”, “communication”, etc.; Fig. S2), suggesting that these more complex functions lie at the intersection of the two systems.

How covarying patterns of gene expression and functional activation emerge over the course of ontogeny and phylogeny remains an open challenge. Patterns of gene expression are involved in cortical reorganization during neurodevelopment, including folding [1], pruning [83] and establishment of cortico-cortical connectivity [74]. In the BrainSpan dataset, we find that the gene-cognition signature has a protracted trajectory over development, gradually becoming most prominent in adulthood (Fig. 5). This external confirmation of our results suggests continued refinement and differentiation of cognitive-affective processes during maturation, but more research is necessary to understand the behavioural consequences of this process at the individual participant level. A related question is how the association between transcription and functional activation evolves across phylogeny. In particular, evolutionary expansion of the cortical mantle is thought to have altered the relationship between molecular gradients and microcircuitry, promoting increasingly complex cognitive function [13, 44]. Thus, the present work could be extended by comparing psychologically-relevant expression patterns, biological processes and cell-type composition across species.

We map whole-genome transcription patterns to a spectrum of cognitive and affective functions across multiple brain areas, but the relationship between gene expression and behaviour has been previously approached from different directions. One approach is to focus on a region of interest. For instance, Vogel and colleagues related variations in cognitive function to a transcriptional gradient across the long axis of the hippocampus [77]. An alternative approach is to map single functions of interest to single genes or gene modules. For instance, Fox and colleagues used Neurosynth and AHBA to identify multiple gene-cognition associations in sub-cortex, including previously established associations between dopamine receptor genes and reward functions in the basal ganglia, as well as novel associations [27]. The results reported here open new possibilities for mapping high-dimensional transcriptional readouts to neurocognitive function in a data-driven and multivariate analysis framework [59], broadening the scope of inquiry to multiple gene sets and comprehensive neurocognitive profiles.

The present work should be understood alongside some important methodological considerations. First, the main analysis involved two singular datasets, potentially limiting the generalizability of the results. Despite extensive validation, the present findings are based on small samples of post-mortem brains and more comprehensive microarray gene expression datasets are necessary for future studies. Second, all analyses were performed in the left cerebral cortices of the six donors, precluding any tests of lateralized brain function, such as language. Third, due to well-documented differences in transcriptional signatures of cortex, subcortex and cerebellum [61], the present investigation focused only on the cortex. How gene expression and functional coactivation covary in subcortical structures should be investigated in future work [27]. Fourth, the mapping of functional activation to psychological terms in Neurosynth cannot distinguish activations from deactivations [86]. Thus, the present results identify gene assemblies whose expression covaries with functional activity, but do not isolate the direction of effect.

In summary, we demonstrate that patterns of gene expression influence cognition and emotion. Organized across a spatially ordered ventromedial-dorsolateral gradient, we show that this genetic signature shapes the composition of cell types and microstructure, ultimately manifesting as a large-scale axis differentiating affective and cognitive processes. Collectively, these results highlight a direct link between molecular dynamics and psychological function.

## METHODS

Code used to conduct the reported analyses are available at https://github.com/netneurolab.

### Microarray gene expression

Regional microarray expression data were obtained from six post-mortem brains provided by the Allen Human Brain Atlas (http://human.brain-map.org/) [40]. Since only two of the six brains included samples from the right hemisphere, analyses were conducted on the left hemisphere only. All processing was performed using the *abagen* toolbox (https://github.com/netneurolab/abagen). These data were processed and mapped to parcellated brain regions at three increasingly finer resolutions, from 34 to 111 left hemisphere cortical grey matter nodes according to the Lausanne anatomical atlas [18, 23].

Microarray probes were reannotated using data provided by Arnatkevičiūtė et al. [3]. A single microarray probe with the highest differential stability, Δ_*S*_ (*p*), was selected to represent each gene [39], where differential stability was calculated as:

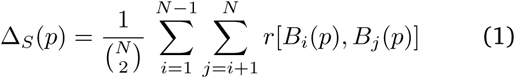

Here, *r* is Spearman’s rank correlation of the expression of a single probe *p* across regions in two donor brains, *B*_*i*_ and *B*_*j*_, and *N* is the total number of donor brains. This procedure retained 20 232 probes, each representing a unique gene.

Next, samples were assigned to brain regions using their corrected MNI coordinates (https://github.com/chrisfilo/alleninf) by finding the nearest region, up to 2mm away. To reduce the potential for misassignment, sample-to-region matching was constrained by hemisphere and cortical/subcortical divisions [3]. If a brain region was not assigned any sample based on the above procedure, the sample closest to the centroid of that region was selected in order to ensure that all brain regions were assigned a value.

Tissue sample expression values were then normalized separately for each donor across genes using a scaled robust sigmoid function [32]:

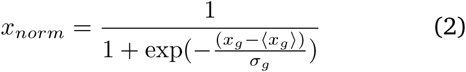

where ⟨*x*_*g*_ ⟩is the median and *σ*_*g*_ is the standard deviation of the expression value of a single gene across regions. Normalized gene expression values were then rescaled to a unit interval:

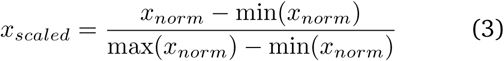

Gene expression values were normalized across tissue samples using the same procedure. Samples assigned to the same brain region were then averaged separately for each donor. Scaled regional expression profiles were finally averaged across donors, resulting in a single matrix **X** with *r* rows corresponding to brain regions and *g* columns corresponding to the retained 20 232 genes. Due to the variability of gene expression across donors, a threshold of 0.1 was imposed on the differential stability of each gene, such that only stable genes were retained for future analysis. At the 34-node, 57-node, and 111-node resolutions, the ensuing number of stable genes retained was 11 560, 10 453, and 8 825, respectively.

### Functional activation

Probabilistic measures of the association between voxels and terms were obtained from Neurosynth, a meta-analytic tool that synthesizes results from more than 15 000 published fMRI studies by searching for high-frequency key words (such as “pain” and “attention”) that are published alongside fMRI voxel co-ordinates (https://github.com/neurosynth/neurosynth [86]). This measure of association is the probability that a given term is reported in the study if there is activation observed at a given voxel. Note that the tool does not distinguish between areas that are activated or deactivated in relation to the term of interest, nor the degree of activation, only that certain brain areas are frequently mentioned in conjunction with certain words. Although more than a thousand terms are reported in Neurosynth, we focus primarily on cognitive function and therefore limit the terms of interest to cognitive and behavioural terms. These terms were selected from the Cognitive Atlas, a public ontology of cognitive science [63]. We used *t* = 123 terms, ranging from umbrella terms (“attention”, “emotion”) to specific cognitive processes (“visual attention”, “episodic memory”), behaviours (“eating”, “sleep”), and emotional states (“fear”, “anxiety”). The coordinates reported by Neurosynth were parcellated into 111 left-hemisphere cortical regions. The probabilistic measure reported by Neurosynth can be interpreted as a quantitative representation of how regional fluctuations in activity are related to psychological processes. For simplicity, we refer to these probabilities as “functional activations” throughout the present report. The full list of terms is shown in Table S1.

### Partial least squares analysis

Partial least squares analysis (PLS) was used to relate gene expression to functional activation. PLS is an unsupervised multivariate statistical technique that decomposes relationships between two datasets (in our case, gene expression, **X**_*n × g*_ and functional activation, **Y**_*n× t*_) into orthogonal sets of latent variables with maximum covariance, which are linear combinations of the original data. This was done by applying singular value decomposition on the matrix **Y**^′^**X** such that

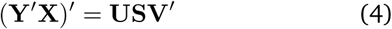

where **U**_*g×t*_ and **V**_*t×t*_ are orthonormal matrices consisting of left and right singular vectors, and **S**_*t*_ *×* _*t*_ is a diagonal matrix of singular values (Fig. 1a) [25]. The *i*^th^ columns of **U** and **V** constitute a latent variable, and the *i*^th^ singular value in **S** represents the covariance between singular vectors. The *i*^th^ singular value is proportional to the amount of covariance between gene expression and functional activation captured by the *i*^th^ latent variable, where the effect size can be estimated as the ratio of the squared singular value to the sum of all squared singular values. In the present study, the left singular vectors (i.e. the columns of **U**) represent the degree to which each gene contributes to the latent variable and demonstrates the extracted association between gene expression and cognitive activation (“gene weights”). The right singular vectors (i.e. the columns of **V**) represent the degree to which the cognitive terms contribute to the same latent variable (“term weights). Positively weighed genes covary with positively weighed terms, and negatively weighed genes covary with negatively weighed terms. Gene and term scores at each brain region for each latent variable can be computed by projecting the original data onto the singular vector weights (Fig. 1c). Positively scored brain regions are regions that demonstrate the covariance between expression of positively weighted genes and activation of positively weighted cognitive terms (and vice versa for negatively scored brain regions):

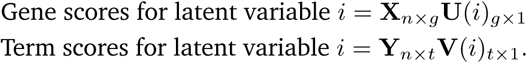

The robustness of the PLS model was assessed by cross-validating the correlation between gene scores and term scores. Since our observations are brain areas and therefore nonindependent, we designed the cross-validation such that the training and testing set were composed of spatially distant brain regions. To achieve this, a random source node and the 75% of brain regions closest in Euclidean distance composed the training set, and the remaining 25% of brain regions composed the testing set (Fig. 1d). PLS was used to compute gene scores and term scores from the training set, as well as the correlation between the two (Corr(**X**_train_**U**_train_, **Y**_train_**V**_train_)). The test set was projected onto the training-derived singular vector weights to generate predicted gene and term scores, and the correlation between predicted scores was computed (Corr(**X**_test_**U**_train_, **Y**_test_**V**_train_)). This procedure was repeated 100 times, yielding a distribution of score correlations for the training and testing sets (Fig. 1d).

### Null model

Spatial autocorrelation-preserving permutation tests were used to assess statistical significance of associations across brain regions, termed “spin tests” [2]. We created a surface-based representation of the parcellation on the FreeSurfer fsaverage surface, via files from the Connectome Mapper toolkit (https://github.com/LTS5/cmp). We used the spherical projection of the fsaverage surface to define spatial coordinates for each parcel by selecting the vertex closest to the center of the mass of each parcel [75]. These vertices were then projected to a sphere, randomly rotated, and reassigned to the closest parcel (10 000 repetitions). The procedure was performed at the parcel resolution rather than the vertex resolution to avoid upsampling the data.

### Gene set analysis

To determine the biological processes in which the gene sets identified by PLS are most involved, we adapted analyses from the Gene Set Enrichment Analysis toolbox (originally available at https://github.com/benfulcher/GeneSetEnrichmentAnalysis [31]). Gene annotations were provided by the Gene Ontology (geneontology.org) and organized such that each biological process category was linked to its associated genes [5, 20]. For each category, we define the category score as the mean loading of the genes of interest, which was done separately for positive genes (genes with the top 50% positive loadings) and negative genes (genes with the top 50% negative loadings). To assess the significance of the category scores, we permuted the rows (brain areas) of the functional activation matrix while preserving spatial autocorrelation using the spherical projection and rotation procedure (spins) described in the previous subsection. We then subjected the original gene expression matrix and the permuted functional activation matrix to PLS and recomputed the category scores (1 000 repetitions).

Next, cell-type deconvolution was performed using cell-specific aggregate gene sets across five human adult postmortem single-cell and single-nucleus RNA sequencing studies ([22, 38, 47, 49, 55, 88]), as presented previously (Supplementary Table 5 from [67]). Briefly, cortical cell classes were determined based on hierarchical clustering of regional topographies across all study-specific cell types in the Allen Human Brain Atlas, resulting in seven major canonical cortical cell classes: astrocytes (Astro), endothelial (Endo), microglia (Micro), excitatory neurons (Neuro-Ex), inhibitory neurons (NeuroIn), oligodendrocytes (Oligo), and oligodendrocyte precursors (OPC) (See Figure 2 from [67]).

### Human Connectome Project dataset

Data from the Human Connectome Project (HCP, S1200 release) [35, 72] was used for measures of cortical thickness, T1w/T2w ratios, functional connectivity, and behavioural tests. The 417 unrelated subjects (age range 22–37 years) with available resting-state fMRI data had individual measures of cortical thickness and T1w/T2w ratios. These structural modalities were acquired on a Siemens Skyra 3T scanner, and included a T1-weighted MPRAGE sequence at an isotropic resolution of 0.7mm, and a T2-weighted SPACE also at an isotropic resolution of 0.7mm. Details on imaging protocols and procedures are available at http://protocols.humanconnectome.org/HCP/3T/imaging-protocols.html. Image processing includes correcting for gradient distortion caused by non-linearities, correcting for bias field distortions, and registering the images to a standard reference space. Measures of cortical thickness are estimated as the geometric distance between the white and grey matter surfaces, and intracortical myelin as the T1w/T2w ratio. Cortical thickness and T1w/T2w ratios for each subject was made available in the surface-based CIFTI file format and parcellated into 219 cortical regions according to the Lausanne anatomical atlas [18]. Only the left-hemisphere regions were retained for analysis.

A group-averaged dense functional connectivity matrix was constructed from the 1 003 subjects with all four 15-minute resting-state fMRI runs. For details on how the dense functional connectivity matrix was constructed, see https://www.humanconnectome.org/storage/app/media/documentation/s1200/HCP1200-DenseConnectome+PTN+Appendix-July2017.pdf. The cortical subset of the matrix was parcellated into 219 nodes according to the Lausanne anatomical atlas [18]. Following Margulies and colleagues, a principal functional gradient was computed by applying diffusion map embedding to the functional connectivity matrix [52], using the Dimensionality Reduction Toolbox (https://lvdmaaten.github.io/drtoolbox/). The procedure yielded an eigenvector map representing the differentiation of unimodal and transmodal cortical regions. Only the left hemisphere was retained for comparison with gene and term scores.

Behavioural performance was assessed using 58 tests. Tests included task performance on sensorimotor processing, cognitive processing, subjective measures of quality of life and personality, and four in-scanner tasks.

When available, age-adjusted scores were used rather than age-unadjusted scores. Demographic variable were excluded. A complete list of behavioural tests used can be found in Table S3. To rule out confounding contributions from genetic correlations among individuals due to family structure, we restricted all behavioural analyses to a subset of 417 individuals with no familial relationships, and with complete T1w/T2w ratio and behavioural performance on the 58 tests.

### External validation using BrainMap

BrainMap is a manually created and curated data repository of results from published functional and structural neuroimaging studies [28, 29, 46, 73]. Specifically, BrainMap includes the brain coordinates that are significantly activated during thousands of different experiments. All experiments conducted on unhealthy subjects were excluded, as well as all experiments without a defined behavioural domain. This resulted in 8 703 experiments organized into 66 unique behavioural domains (Table S2). To enable more direct comparability with results using Neurosynth, the BrainMap data were subjected to the Neurosynth meta-analytic pipeline (https://github.com/neurosynth/neurosynth). This approach resulted in a region by term matrix of probabilistic measures that certain terms are published in conjunction with certain brain regions.

### External validation using BrainSpan

BrainSpan is a database of gene expression in the brain across development, available at https://www.brainspan.org/static/download.html [58]. Gene expression levels were quantified in specific tissue samples from post-mortem brains ranging from eight post-conception weeks (pcw) to 40 years of age. Ages were binned into five life stages: fetus (8pcw–37pcw), infant (4mos–1yr), child (2yrs–8yrs), adolescent (11yrs–19yrs), and adult (21yrs–40yrs) [82]. For each age category, a gene expression matrix was constructed by averaging the expression of every gene across identical regions. Any missing data was replaced with the median expression of the gene across all regions. Of the sixteen unique cortical brain regions with gene expression levels, four regions did not have gene expression estimates in any of the age categories besides the fetal stage. For comparison with PLS results derived from AHBA, we selected the 9 568 available genes with differential stability greater than 0.1 as defined on the 34-node parcellation. Gene scores for the 12 regions with expression levels available for all age stages were estimated by projecting the gene expression matrices onto the PLS-derived gene weights.

To relate estimated gene scores with term scores, we defined a region-to-region correspondence map from the 16-node parcellation to the 34-node parcellation. Term scores were averaged across sibling nodes such that a single term score was available for all 16 regions available in BrainSpan. Note that brain regions in BrainSpan are not organized by hemisphere; therefore, regional expression levels in the 16 regions are not necessarily measured from the left hemisphere only.

## Acknowledgments

We thank Vincent Bazinet, Laura Suarez, Golia Shafiei, Bertha Vazquez-Rodriguez and Zhen-Qi Liu for their comments and suggestions on the manuscript. This research was undertaken thanks in part to funding from the Canada First Research Excellence Fund, awarded to McGill University for the Healthy Brains for Healthy Lives initiative. BM acknowledges support from the Natural Sciences and Engineering Research Council of Canada (NSERC Discovery Grant RGPIN #017-04265) and from the Canada Research Chairs Program. DB was supported by the Healthy Brains Healthy Lives initiative (Canada First Research Excellence fund), the CIFAR Artificial Intelligence Chairs program (Canada Institute for Advanced Research), Google (Research Award), and by NIH grant R01AG068563A.

---

**Figure S1.**
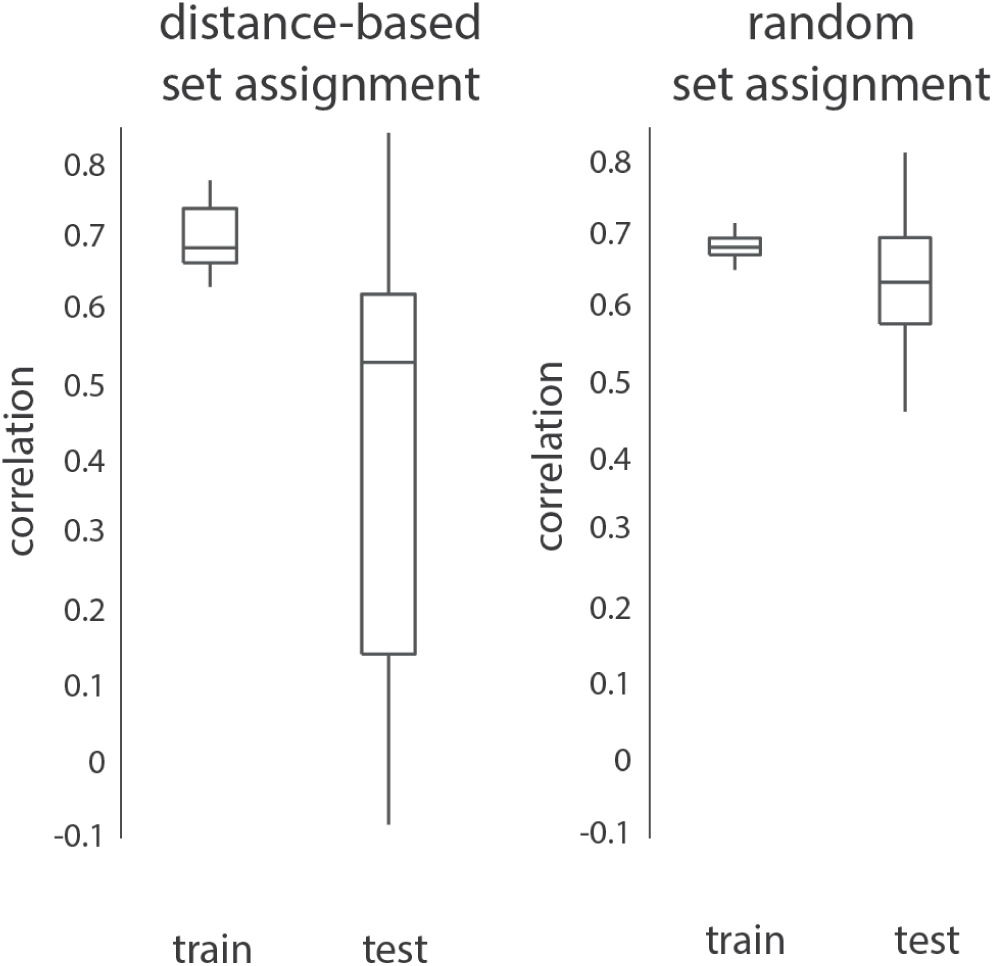
Cross-validation using distance-based set assignment results in more conservative score correlations. Left: In distance-based set assignment, the 75% of nodes closest to a randomly chosen source node are assigned to the training set, and the remaining 25% of nodes are assigned to the testing set. Right: In random set assignment, training and test sets are assigned randomly. Due to spatial autocorrelation, random assignment yields inflated out-of-sample performance estimates. Only distance-based assignment was used in the manuscript. Random set assignment is shown only for comparison.

**TABLE S1.**
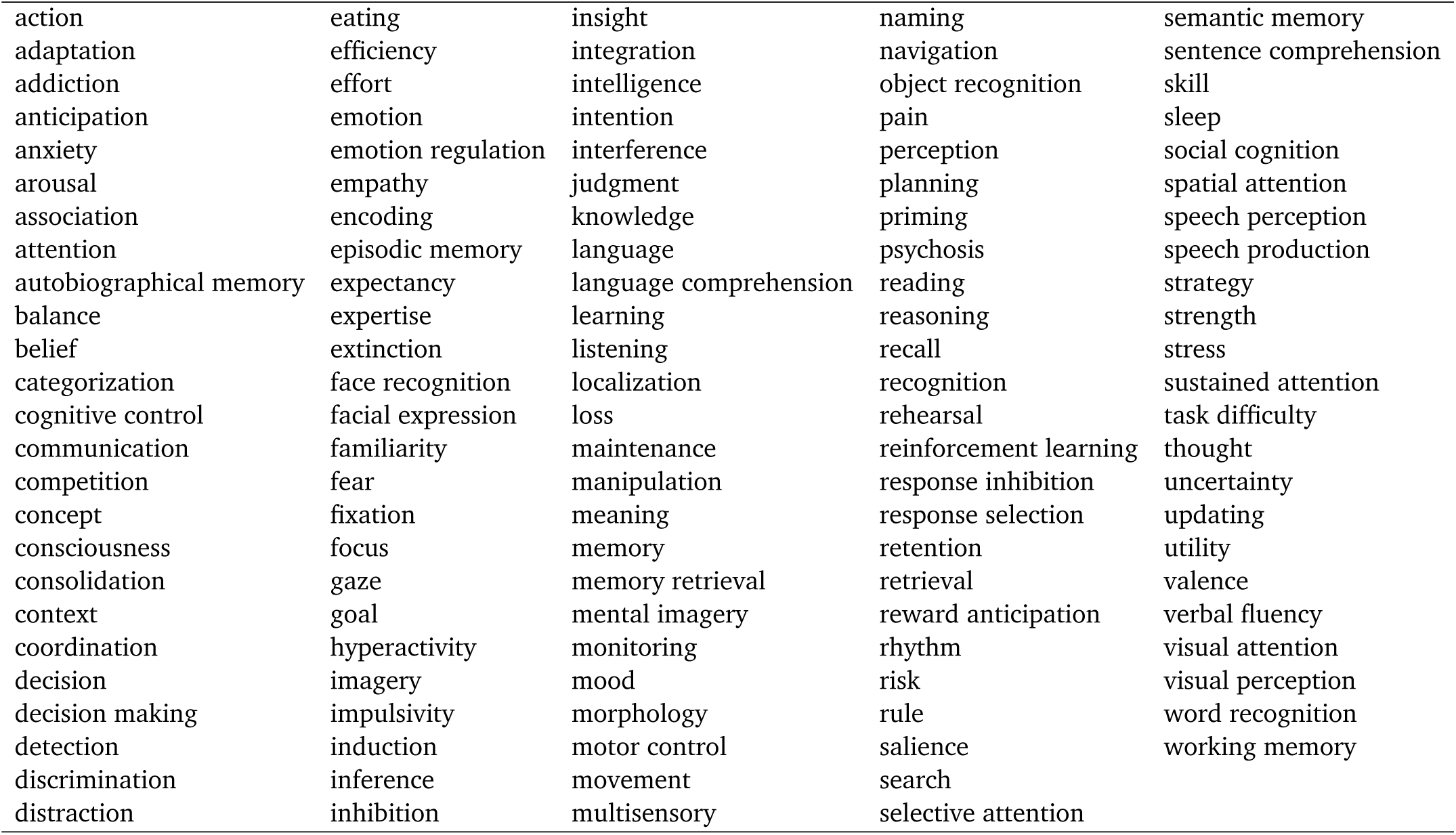
Neurosynth terms. Terms that overlapped between the Neurosynth database [86] and the Cognitive Atlas [63] were included in analyses.

|  |  |  |  |  |
| --- | --- | --- | --- | --- |
| action | eating | insight | naming | semantic memory |
| adaptation | efficiency | integration | navigation | sentence comprehension |
| addiction | effort | intelligence | object recognition | skill |
| anticipation | emotion | intention | pain | sleep |
| anxiety | emotion regulation | interference | perception | social cognition |
| arousal | empathy | judgment | planning | spatial attention |
| association | encoding | knowledge | priming | speech perception |
| attention | episodic memory | language | psychosis | speech production |
| autobiographical memory | expectancy | language comprehension | reading | strategy |
| balance | expertise | learning | reasoning | strength |
| belief | extinction | listening | recall | stress |
| categorization | face recognition | localization | recognition | sustained attention |
| cognitive control | facial expression | loss | rehearsal | task difficulty |
| communication | familiarity | maintenance | reinforcement learning | thought |
| competition | fear | manipulation | response inhibition | uncertainty |
| concept | fixation | meaning | response selection | updating |
| consciousness | focus | memory | retention | utility |
| consolidation | gaze | memory retrieval | retrieval | valence |
| context | goal | mental imagery | reward anticipation | verbal fluency |
| coordination | hyperactivity | monitoring | rhythm | visual attention |
| decision | imagery | mood | risk | visual perception |
| decision making | impulsivity | morphology | rule | word recognition |
| detection | induction | motor control | salience | working memory |
| discrimination | inference | movement | search |  |
| distraction | inhibition | multisensory | selective attention |  |

**Figure S2.**
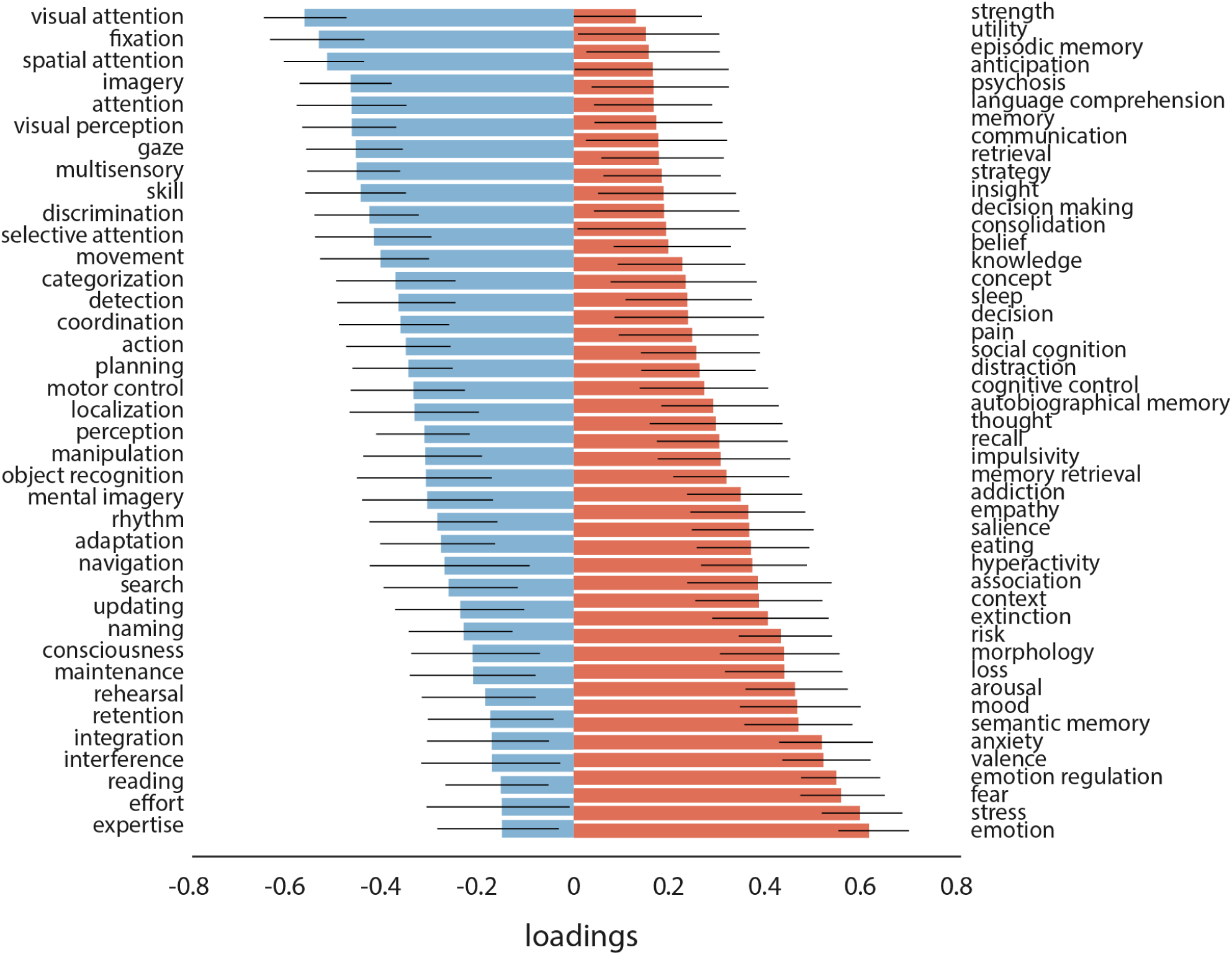
Neurosynth term loadings. The loading for each term is calculated as the correlation between functional activation across brain regions and PLS-derived gene scores. Error bars indicate bootstrap-estimated 95% confidence intervals. All terms with a confidence interval that changes sign are excluded.

**TABLE S2.** BrainMap terms. BainMap terms are organized by behavioural domain. All 66 unique behavioural domain (excluding any undefined domains) used in analyses are shown here.

|  |  |  |  |  |
| --- | --- | --- | --- | --- |
| air-hunger | disgust | language | phonology | speech (action) |
| alcohol | emotion | learning | preparation | speech (languag) |
| amphetamines | estrogen | marijuana | psychiatric medications | SSRIs |
| anger | execution | memory | reasoning | steroids and hormones |
| antidepressants | explicit | motion | rest | syntax |
| antipsychotics | fear | music | sadness | thermoregulation |
| anxiety | gustation | nicotine | semantics | thirst |
| attention | happiness | non-steroidal anti-inflammatory drugs | sexuality | time |
| audition | humour | observation | shape | vision |
| bladder | hunger | olfaction | sleep | working memory |
| caffeine | imagination | opioids | social cognition |  |
| capsaicin | inhibition | orthography | soma |  |
| cognition | interoception | pain | somesthesis |  |
| colour | ketamine | pharmacology | space |  |

**Figure S3.**
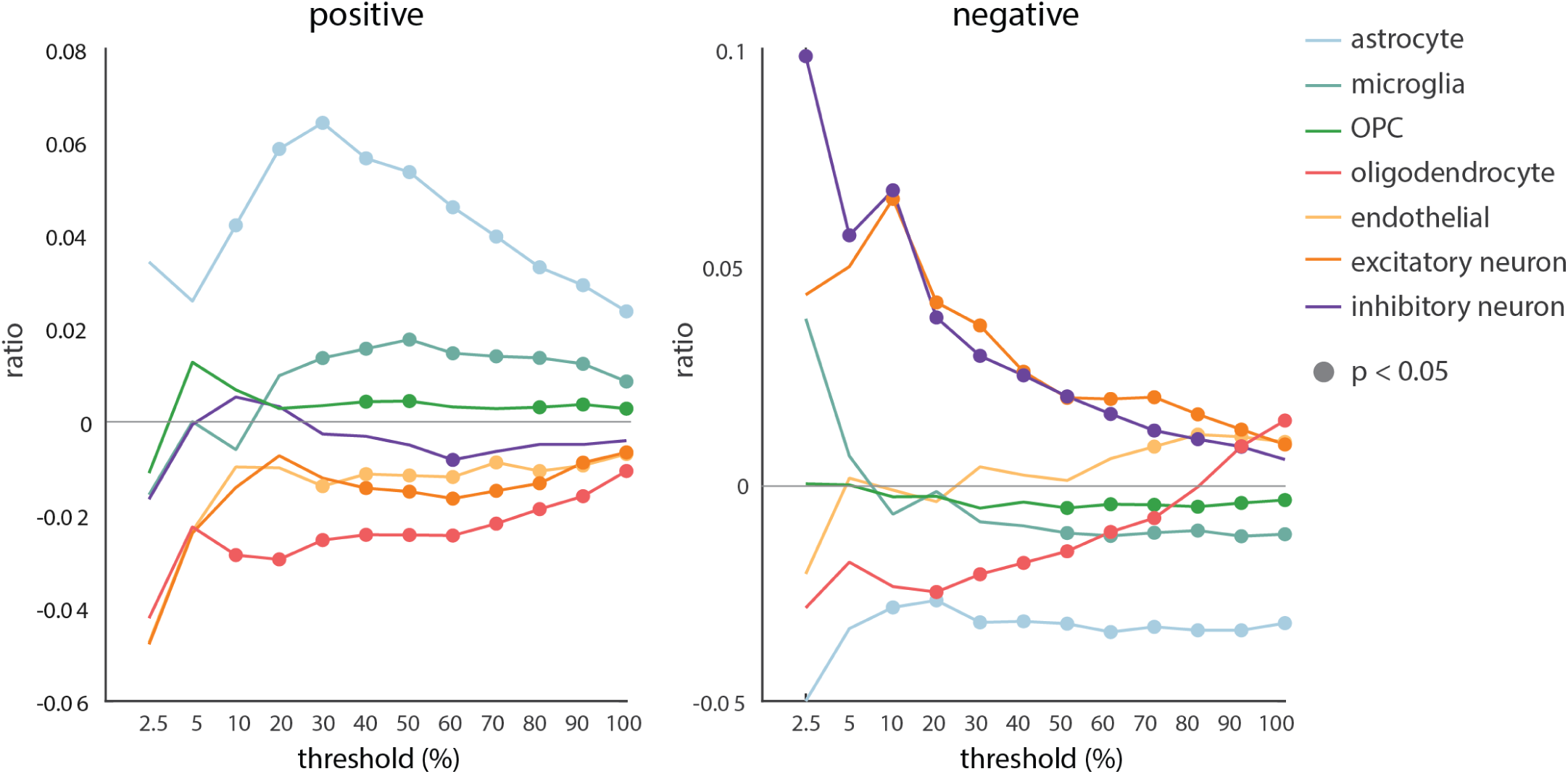
Specific cell-type expression is consistent across gene sets. Positive and negative gene sets were constructed using the largest positive/negative loadings, ranging from the top 2.5% genes to all genes. Each curve represents the difference between the ratio of genes in each gene set preferentially expressed in a cell-type and the mean null ratio, computed from random gene sets (10 000 repetitions). Curves above zero indicate overexpression and curves below zero indicate underexpression. Circles demonstrate significance.

**Figure S4.**
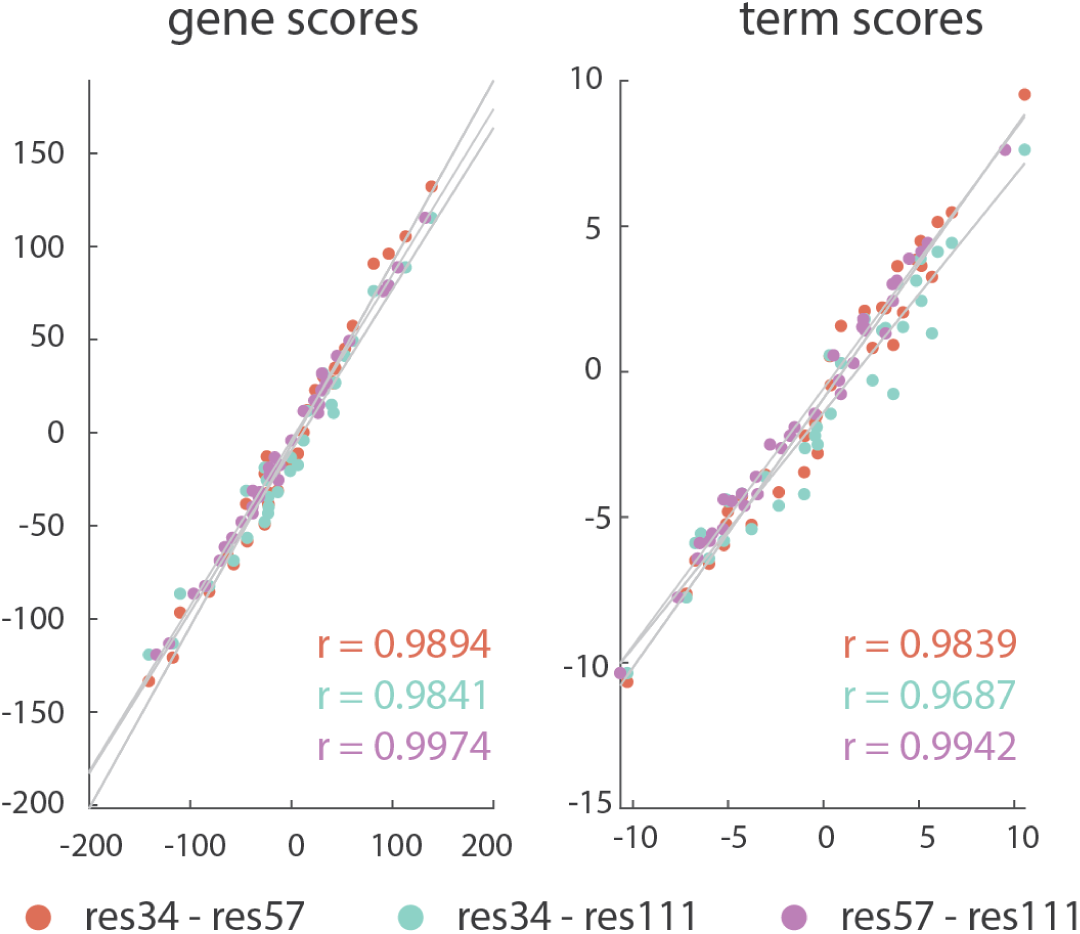
Gene and term scores are consistent across parcellation resolutions. PLS was performed on gene expression and functional activation matrices at three progressively finer parcellation resolutions (*n* = 34, *n* = 57, and *n* = 111 left hemisphere cortical regions; [18]). The resulting gene and term scores at each resolution were then correlated with the gene and term scores from other resolutions.

**Figure S5.**
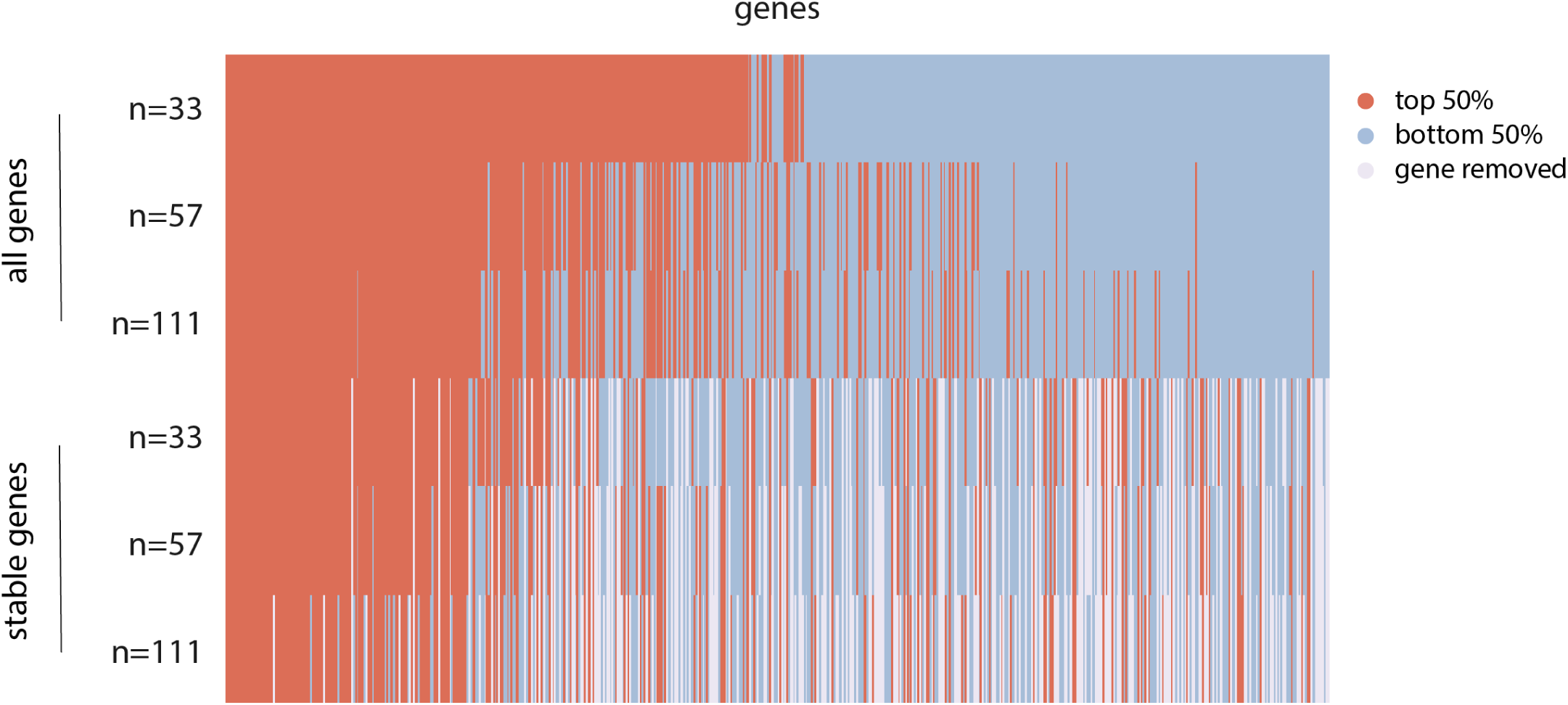
Reliable gene sets are consistent across parcellation resolutions and gene set assignment. The reliability of each gene, as defined by its loading, was recomputed for six different gene expression matrices (3 parcellation resolutions *×* 2 gene set assignment strategies). Tuning the parcellation of brain regions (*n* = 34, *n* = 57, and *n* = 111 left hemisphere cortical regions) and the set of genes (all genes or differentially stable genes) used in the gene expression matrix reveals reliable gene sets are consistent across different methodological choices when constructing the gene expression matrix. Reliable genes are coloured red (top 50% of positive/negative loadings), unreliable genes are blue (bottom 50% of positive/negative loadings), and genes removed from the analysis because their differential stability is less than 0.1 are white.

**Figure S6.**
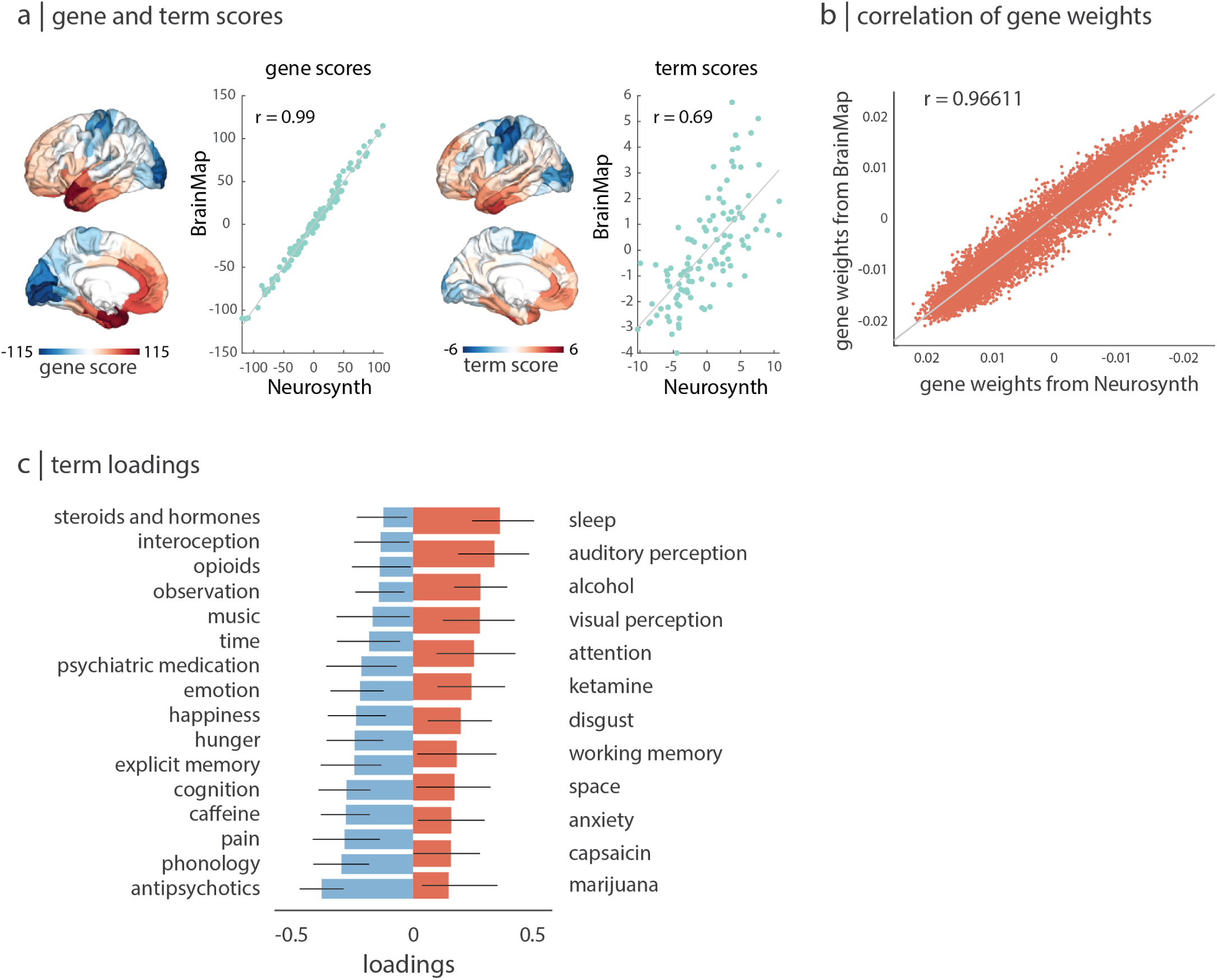
Replication using BrainMap. Gene expression was related to a functional activation matrix derived using the manually-curated BrainMap [28, 29]. (a) PLS-derived gene and term scores are correlated between Neurosynth- and BrainMap-derived functional activation matrices. (b) PLS-estimated gene weights for the first latent variable from the Neurosynth- and BrainMap-derived functional activation matrix are correlated. (c) Reliable terms with positive loadings (red) and negative loadings (blue). Error bars indicate bootstrap-estimated 95% confidence intervals.

**TABLE S3.** HCP behavioural measures. Behavioural measures that were included in analyses cover a range of cognitive and emotional tasks. Demographic variables were excluded.

|  |  |
| --- | --- |
| Pittsburg Sleep Quality Index Score | Crystallized Cognition Composite Score (age adjusted) |
| Picture Sequence Memory Test (age adjusted) | Penn Emotion Recognition (correct responses) |
| Dimensional Change Card Sort Test (age adjusted) | Penn Emotion Recognition (reaction time) |
| Flanker (age adjusted) | Penn Emotion Recognition (anger, correct responses) |
| Penn Matrix Test (correct response) | Penn Emotion Recognition (fear, correct responses) |
| Penn Matrix Test (skipped items) | Penn Emotion Recognition (happiness, correct responses) |
| Penn Matrix Test (reaction time) | Penn Emotion Recognition (neutral, correct responses) |
| Reading Test (age adjusted) | Penn Emotion Recognition (sadness, correct responses) |
| Picture Vocabulary Test (age adjusted) | anger |
| Pattern Comparison Processing Test (age adjusted) | hostility |
| Delay Discounting (AUC \$200) | aggression |
| Delay Discounting (AUC \$40 000) | fear |
| Penn Line Orientation (total correct) | anxiety |
| Penn Line Orientation (reaction time) | sadness |
| Penn Line Orientation (incorrect) | life satisfaction |
| Short Penn Continuous Performance Test (true positives) | life purpose |
| Short Penn Continuous Performance Test (true negatives) | positive affect |
| Short Penn Continuous Performance Test (false positives) | friendship |
| Short Penn Continuous Performance Test (false negatives) | loneliness |
| Short Penn Continuous Performance Test (reaction time) | perceived hostility |
| Short Penn Continuous Performance Test (sensitivity) | perceived rejection |
| Short Penn Continuous Performance Test (specificity) | emotional support |
| Short Penn Continuous Performance Test (longest run of non-responses) | instrumental support |
| Penn Word Memory (total correct) | stress |
| Penn Word Memory (reaction time) | self-efficacy |
| List Sort (age adjusted) | in-scanner emotion task |
| Fluid Cognition Composite Score (age adjusted) | in-scanner language task |
| Early Childhood Composite Score (age adjusted) | in-scanner relational task |
| Cognitive Function Composite Score (age adjusted) | in-scanner working memory task |

